# CCorGsDB: A Database for Clock Correlated Genes in the Mouse and Human Central Nervous Systems

**DOI:** 10.1101/2025.06.11.659048

**Authors:** José Luiz Araújo Santos, Vinícius Tenório Braga Cavalcante Pinto, Thales Eduardo da Silva Santos, Daniel Gomes Coimbra, Tiago Gomes de Andrade

**Affiliations:** Circadian Medicine Center, Faculty of Medicine, Federal University of Alagoas, Maceió, Alagoas, Brazil

**Keywords:** Bioinformatics, Central nervous system, Circadian rhythms, Gene Co-expression Networks, Gene expression

## Abstract

We introduce CCorGsDB, a web-based tool that integrates co-expression networks filtered by circadian biomarkers to identify candidate clock-regulated genes in spatially defined regions of the mouse and human central nervous systems. Built using Weighted Gene Co-expression Network Analysis (WGCNA), CCorGsDB includes ∼16,000 genes for mouse and ∼37,000 for human, highlighting genes highly correlated with canonical clock markers. The tool incorporates disease associations and drug target information for compounds with short half-lives acting on the CNS, supporting chronopharmacological research. Users can explore region-specific data through an interactive interface offering query, visualization, and download options. CCorGsDB is freely accessible at https://famed.ufal.br/ccorgs.

---

Circadian rhythms regulate diverse biological processes such as cell division, hormone secretion, sleep, cognition, and mood [1]. In mammals, these rhythms are governed by transcriptional-translational feedback loops involving clock genes and their downstream effectors, the clock-controlled genes (CCGs) [2]. Up to 50% of the mammalian transcriptome exhibits rhythmicity [3], and many CCGs are drug targets with therapeutic relevance [4].

Although the suprachiasmatic nucleus (SCN) is the master circadian pacemaker [5], many brain regions display rhythmic activity independent of the SCN [6], and circadian disruption is a hallmark of several neuropsychiatric disorders [7]. Identifying region-specific CCGs in the central nervous system (CNS) is thus critical for understanding circadian regulation in brain physiology and pathology [8].

Clock genes are broadly expressed [9], but CCGs exhibit high tissue specificity [10], complicating their identification in the CNS, particularly in humans, where time-series transcriptomic data are limited [3,11–19]. Moreover, not all circadian-relevant genes show rhythmic expression; post-transcriptional regulation can decouple transcript and protein rhythms [20–23].

To address these limitations, we constructed CCorGsDB, a database of Clock Correlated Genes (CCorGs) derived from WGCNA-based co-expression networks in mouse and human CNS regions, filtered by their correlation with ten canonical clock genes. This approach captures potential circadian regulators beyond cycling transcripts. Details on data preprocessing, network construction, and gene filtering are provided in the Supplementary Material.

Compared to generic co-expression resources, CCorGsDB integrates circadian biomarkers to refine gene selection, incorporating ∼16k (mouse) and ∼37k (human) genes. The database also links CCorGs to neurodegenerative and behavioral disorders, and to CNS-active drugs with short half-lives (≤12 h), highlighting its utility in chronopharmacology [24–26]. CCorGsDB is publicly available at https://famed.ufal.br/ccorgs.

We validated the database using time-series datasets from sixteen mouse CNS regions [3,17]. Genes with higher circadian amplitudes (rAMP) consistently showed stronger correlations in CCorGsDB, supporting the circadian relevance of these networks (Supplementary Table 2, Figure 1A–B). An exception was the lateral hypothalamus-rostral region, which approached but did not reach statistical significance (p = 0.05259). Clock gene enrichment was also significant in multiple networks for both species (Figure 1C, Supplementary Table 3). Notably, Per3 was the top-ranked CCorG in the human cerebellum (r = 0.847, p = 9.9e–68), while Hnrnpdl and Akt3 were the strongest correlates in mouse CNS (r = 0.903) and human hypothalamus (r = 0.919), respectively—both with known links to circadian function [27,28].

**Figure 1.**
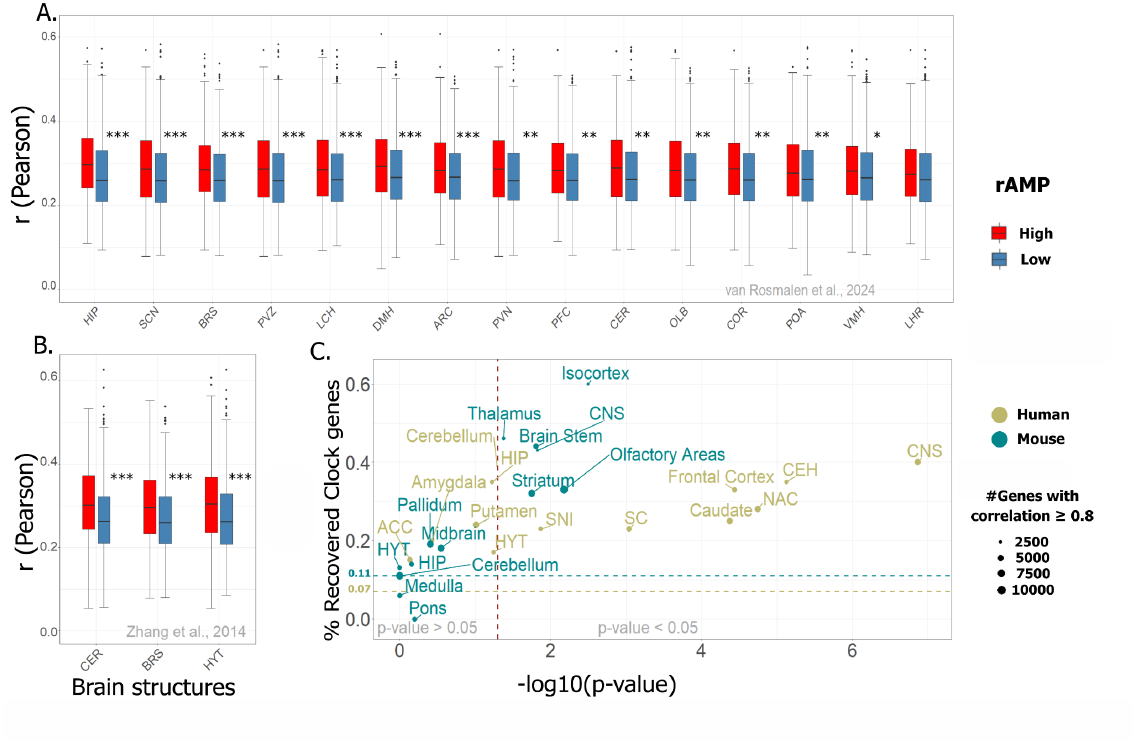
Clock correlated genes (CCorGs) are enriched with both cycling transcripts and clock genes. Correlation values were higher in the subset of genes with higher amplitudes (95th percentile) compared to those with lower amplitudes (5th percentile) across all analyzed tissues, except in the LHR (MW, p = 0.05259). Boxplots indicate median and 95% confidence intervals. Time course experiments were obtained from van Rosmalen et al., 2024 (A) and Zhang et al., 2014 (B). *** indicates p ≤ 0.001; ** p ≤ 0.01; * p < 0.05. C. The ratio of clock genes recovered in the 90th percentile of CCorGs (with Fdr < 0.05) are higher in 13 regions compared to the mean ratio in the input samples for mice (0.11) and humans (0.07) (Fisher’s Exact Test p < 0.05). Vertical dashed line indicates the p.value cutoff of 0.05. Horizontal dashed lines indicate average clock genes ratio in mouse and human input datasets for WGCNA. Supplementary Tables 2 and 3 detail the statistical parameters. relative Amplitude (rAMP); central nervous system (CNS); cerebellar hemisphere (CEH); spinal cord (SC); substantia nigra (SNI); nucleus accumbens (NAC); anterior cingulate cortex (ACC); arcuate nucleus of hypothalamus (ARC); brainstem (BRS); cerebellum (CER); whole cortex (COR); prefrontal cortex (PFC); olfactory bulb (OLB); hippocampus (HIP); dorsomedial hypothalamus (DMH); preoptic area (POA); suprachiasmatic nuclei (SCN); paraventricular nuclei of hypothalamus (PVN); lateral hypothalamus-caudal region (LCH); lateral hypothalamus-rostral region (LHR); ventromedial hypothalamus (VMH); periventricular zone (PVZ); whole hypothalamus (HYT).

CCorGsDB also revealed hundreds of genes associated with neurodegenerative and behavioral disorders, as identified via DisGeNET [32]. Some are targets of antiepileptic and dopaminergic drugs whose efficacy improves with time-specific administration [33–35], reinforcing the value of CCorGsDB for chronotherapeutic strategies.

In preclinical settings, the database can assist in selecting genes for functional assays or testing drug timing in disease models. Clinically, it may support biomarker discovery and therapeutic optimization. Future integration with machine learning could refine predictions and stratifications of circadian phenotypes.

## Conclusion

Our approach enables the identification of candidate circadian genes across CNS regions, offering a practical alternative where time-series data are lacking. CCorGsDB extends the reach of circadian transcriptomics by capturing functionally relevant genes beyond rhythmic expression. With its web interface, visual tools, and translational features, it serves as a resource for chronobiology, neuroscience, and molecular medicine.

## Acknowledgments

We thank John Hogenesch, Marc Ruben, Gang Wu and Daniel Gitaí for thoughtful discussions, and CNPq, CAPES, and FAPEAL for financial support. The mouse raw data used for the analyses described in this article were obtained from the Allen Mouse Brain Atlas (http://mouse.brain-map.org/) between the years 2017 and 2021. The human data were collected from the Genotype-Tissue Expression (GTEx) portal (https://gtexportal.org/home/) in 2021 and 2022.

