## Supplementary material for "CCorGsDB: A Database for Clock Correlated Genes in the Mouse and Human Central Nervous Systems"

### Methods

The mouse data were obtained from the Allen Mouse Brain Atlas database (http://mouse.brain-map.org/) [34], downloaded between 2017 and 2021. We collected the in-situ hybridization expression data (expression energy) of 15,951 genes (15,804 protein-coding) in thirteen mouse CNS tissues: brain stem, cerebellum, hippocampus, hypothalamus, isocortex, medulla, midbrain, olfactory areas, pallidum, pons, striatum, and thalamus, in both sagittal and coronal anatomical series [35], using the ABAData R package [36]. We additionally unified the data from all tissues into a single dataset, termed the "mouse central nervous system." For each gene in each tissue, expression data were collected from at least 20 common subregions among the total set of genes. Within this set, each gene could have more than one expression data point per sub-region, corresponding to one or more experimental replicas according to the anatomical plane and different gene sequences. The cortical subplate was not included in the analysis because it did not meet the criteria for a minimum number of subregions.

The human data were collected from the Genotype-Tissue Expression (GTEx) portal (https://gtexportal.org/home/) [37], downloaded in 2021 and 2022. We obtained the RNA-sequencing expression data of 37,464 genes (16,328 protein-coding) in twelve distinct human brain subregions of postmortem donors: anterior cingulate cortex, amygdala, caudate, cerebellar hemisphere, cerebellum, frontal cortex, hippocampus, hypothalamus, nucleus accumbens, putamen, spinal cord, and substantia nigra. We considered all samples by brain subregion, estimated at over a hundred for each of them. As was done with mouse data, we unified data from all human tissues into a single dataset, termed the "human central nervous system."

Co-expression networks were built using the 'WGCNA' (Weighted Gene Correlation Network Analysis) package [38]. Twelve signed-type weighted gene co-expression networks were constructed for each species, one for each tissue, and additional networks for datasets integrating all tissues, described here as Central Nervous System (CNS) networks for mice and humans. Hierarchical clustering by average linkage was implemented to detect outliers. Pearson's correlations between each gene pair were calculated to build an adjacency matrix using the "cor" function. A soft-threshold power for each co-expression network was calculated to achieve approximate scale-free topology. Then, the topological overlap measure

(TOM) and corresponding dissimilarity (1-TOM) were calculated using an adjacency matrix. 1-TOM was used as a distance for gene hierarchical clusters, and the Dynamic Tree Cut algorithm and blockwiseModules function were used to identify the modules, defined as clusters of highly interconnected genes according to their similarity of expression profiles [39]. In all networks, the minimum number of genes per module was 30, and a limit of 0.25 was used in the "cutheight" argument to determine the height at which the branches should be cut. In each module, we identified the module eigengene (ME) using the "moduleEigengenes" function, considered as a representative summary of the gene expression profile in a module and the first principal component of a given module [40]. Hub genes were determined by the highest connectivity in candidate modules, measured by Module Membership.

To find relevant gene modules in each co-expression network, we created module–trait relationships based on the correlation between ME and circadian clock traits. We considered traits to be the expression from ten genes recognized as the best biomarkers for circadian rhythms in 12 mouse tissues based on a machine learning algorithm: Arntl, Cry1, Dbp, Npas2, Nr1d1, Nr1d2, Per1, Per2, Per3, and Tef [26]. Eight of these are known to be core components of the molecular clock: Arntl (Bmal1), Cry1, Per1, Per2, Per3, Nr1d1 (Rev-erbα), Nr1d2 (Rev-erbβ), and Npas2 [2]. Dbp and Tef are transcription factors mediating the circadian expression of many downstream genes [41]. Because each circadian gene can have one or more experimental replicas in ABA, twenty-two traits were used in total for each co-expression network construction. A correlation coefficient $(r) \geq 0.8$ and a p-value $< 0.05$ were set as the criteria for significant correlation between a given ME and every circadian clock biomarker. After determining the biologically significant modules, we calculated the gene significance (GS) (the Pearson's correlation coefficient between the gene and the circadian clock biomarkers) [26]. We then bootstrapped the correlations in 1,000 simulations using the "boot" package to estimate the 95% confidence intervals.

We also evaluated different mouse encephalon tissues with available circadian transcriptome datasets, as a strategy to validate CCorGs as candidate genes associated with the circadian clock. From Zhang et al., 2014 [3], we obtained brainstem, cerebellum, and hypothalamus from C57/BL6 mice. From van Rosmalen, 2024 [17], we analyzed the cortex, prefrontal cortex, olfactory bulb, hippocampus, preoptic area, suprachiasmatic nuclei, arcuate nucleus, paraventricular nuclei of hypothalamus, lateral hypothalamus-caudal region, lateral hypothalamus-rostral region, dorsomedial hypothalamus, ventromedial hypothalamus, and hypothalamic periventricular zone. In this case, only data from CBA/CaJ nocturnal mice and from equivalent regions to those present in the CCorGsDB database were used. Data normalization was performed by tissue using the DESeq2 R package [42]. The rAmp (relative amplitude) of each gene was obtained using the MetaCycle R package [43]. We used the Mann-Whitney test to evaluate the difference between Pearson correlation values in the 5th and 95th percentiles of the rAmp distribution in each region. Cohen's d test was used to calculate the effect size. Additionally, we used the Fisher exact test to evaluate the possible enrichment for clock genes in a subset of CCorGs (90th percentile of GS positive correlation values—FDR < 0.05—in significant modules) based on a list of 22 genes (Arntl, Arntl2, Clock, Cry1, Cry2, Csnk1a1, Csnk1d, Csnk1e, Dbp, Fbxl3, Fbxl21, Hlf, Npas2, Nr1d1, Nr1d2, Per1, Per2, Per3, Rora, Rorb, Rorc, and Tef) which are important molecular components for the circadian clock [44, 45]. We considered a 95% confidence interval and an alpha of 0.05 as a cutoff for statistical significance.

Neuropsychiatric disorders, cognitive, and behavioral traits associated with CCorGs were identified based on the Disgenet database [32], using the disgenet2r R package [46]. Pharmacological drugs with a half-life of up to 12 hours acting on the CNS were collected from the DrugBank database (https://go.drugbank.com/) [47]. The database was implemented using PHP, Javascript, HTML, JQuery, MySQL, Plotly.js, Bootstrap, and Ajax, and is available as a public domain website at https://famed.ufal.br/ccorgs. Dynamic filtering of the available datasets is provided based on different statistics and non-statistical parameters. Downloadable results for each search include images in PNG format and CSV files with reported statistics.

**Supplementary Table 1**. Mouse and human tissues, input parameters used for Weigheted Gene Co-expression Network Analysis (WGCNA) and overall positive correlations for CCorGs sets within networks. Mean, SD (standard deviation), rmin (minimum GS Pearson correlation), rmax (maximum GS Pearson correlation).

| Tissues | Number of Samples | Number of Input genes | *Soft-Threshold Power* (β) | $r_{min}$ | $r_{max}$ | Mean | SD |
|---|---|---|---|---|---|---|---|
| **Mouse** | | | | | | | |
| Central Nervous System | 89 | 19933 | 8 | 0.0020 | 0.9036 | 0.55 | 0.17 |
| Isocortex | 62 | 19933 | 6 | 0.0072 | 0.9344 | 0.58 | 0.16 |
| Hippocampus | 33 | 19932 | 7 | 0.0026 | 0.9694 | 0.66 | 0.17 |
| Medulla | 30 | 19931 | 8 | 0.0009 | 0.9867 | 0.68 | 0.15 |
| Midbrain | 22 | 19933 | 8 | 0.0003 | 0.9917 | 0.69 | 0.15 |
| Brain Stem | 22 | 19933 | 7 | 0.0001 | 0.9820 | 0.65 | 0.19 |
| Pallidum | 21 | 19923 | 8 | 0.0074 | 0.9961 | 0.72 | 0.16 |
| Thalamus | 21 | 19924 | 8 | 0.0001 | 0.9617 | 0.56 | 0.18 |
| Olfactory Areas | 20 | 19933 | 10 | 0.0005 | 0.9914 | 0.71 | 0.16 |
| Cerebellum | 20 | 19924 | 9 | 0.0006 | 0.9928 | 0.74 | 0.17 |
| Pons | 20 | 19932 | 8 | 0.0060 | 0.9770 | 0.71 | 0.15 |
| Striatum | 20 | 19933 | 8 | 0.0008 | 0.9916 | 0.69 | 0.17 |
| Hypothalamus | 20 | 19885 | 6 | 0.0034 | 0.9931 | 0.70 | 0.16 |
| **Human** | | | | | | | |
| Central Nervous System | 2642 | 53921 | 6 | 0.0006 | 0.9595 | 0.61 | 0.18 |
| Anterior Cingulate Cortex | 176 | 17958 | 8 | 0.0007 | 0.9552 | 0.67 | 0.17 |
| Amygdala | 152 | 17528 | 12 | 0.0032 | 0.9179 | 0.64 | 0.13 |
| Caudate | 246 | 51311 | 10 | 0.0009 | 0.9659 | 0.58 | 0.19 |
| Cerebellar Hemisphere | 215 | 50943 | 8 | 0.0003 | 0.9331 | 0.56 | 0.17 |
| Cerebellum | 241 | 17526 | 9 | 0.0513 | 0.8476 | 0.58 | 0.14 |
| Frontal Cortex | 209 | 51004 | 8 | 0.0014 | 0.9409 | 0.58 | 0.19 |
| Hippocampus | 197 | 17549 | 8 | 0.0011 | 0.9363 | 0.62 | 0.17 |
| Hypothalamus | 201 | 51222 | 10 | 0.0060 | 0.9188 | 0.59 | 0.17 |
| Nucleus Accumbens | 246 | 51506 | 8 | 0.0003 | 0.9660 | 0.58 | 0.20 |
| Putamen | 205 | 17521 | 10 | 0.0059 | 0.9722 | 0.69 | 0.15 |
| Spinal Cord | 159 | 50064 | 10 | 0.0005 | 0.9253 | 0.55 | 0.22 |
| Substantia Nigra | 139 | 49596 | 8 | 0.0076 | 0.9273 | 0.60 | 0.19 |

**Supplementary Table 2**. Mann-Whitney test was conducted to evaluate the difference between Gene Significance (GS) values in the 5th and 95th percentiles of the rAMP distribution in each region analyzed from Van Rosmalen, et al., 2024 [62] and Zhang et al., 2014. Cohen's d test was used to evaluate the size effect.

| Region | Corresponding area in CCorGsDB | GS mean / median in $5^{th}$ percentile of rAMP (#CCorGS) | GS mean / median in $95^{th}$ percentile of rAMP (#CCorGs) | Mann-Whitney p-value | Size effect (Cohen's d) |
|---|---|---|---|---|---|
| **Van Rosmalen et al., 2024** | | | | | |
| Arcuate nucleus | Hypothalamus | 0.2723037 / 0.2667017 (998) | 0.2928806 / 0.2823499 (361) | p = 0.0002941 | d = 0.809557 |
| Brainstem | Brainstem | 0.267867 / 0.2592915 (998) | 0.293323 / 0.2843633 (326) | p = $3.487e^{-06}$ | d = 1.037497 |
| Cerebellum | Cerebellum | 0.2749704 / 0.2620485 (979) | 0.2897903 / 0.2843633 (280) | p = 0.005329 | d = 0.6230717 |
| Whole cortex | Isocortex | 0.2693477 / 0.2597189 (990) | 0.2903803 / 0.2866571 (250) | p = 0.00608 | d = 0.7665251 |
| Prefrontal cortex | Isocortex | 0.2694226 / 0.2591665 (1087) | 0.288876 / 0.2826808 (338) | p = 0.0004363 | d = 0.5933874 |
| Olfactory Bulb | Olfactory Areas | 0.2693477 / 0.2597189 (990) | 0.2891894 / 0.2826808 (345) | p = 0.005562 | d = 0.7719158 |
| Hippocampus | Hippocampus | 0.2738964 / 0.2593759 (1015) | 0.3034936 / 0.2966472 (413) | p = $2.856e^{-09}$ | d = 1.328156 |
| Dorsomedial Hypothalamus | Hypothalamus | 0.2756436 / 0.2659488 (1000) | 0.2963376 / 0.2919925 (327) | p= 0.0001992 | d = 0.8318283 |
| Preoptic area | Hypothalamus | 0.272095 / 0.2613014 ( (1006) | 0.2872428 / 0.2761689 (338) | p = 0.007576 | d = 0.5971168 |
| Suprachiasmatic Nuclei | Hypothalamus | 0.2702148 / 0.2605448 (962) | 0.2964105 / 0.2955416(384) | p = $3.329^{e-07}$ | d = 1.141245 |
| Paraventricular Nuclei | Hypothalamus | 0.2713585 / 0.2584326 (966) | 0.2908163 / 0.2858588 (381) | p = 0.0004358 | d = 0.7865114 |
| Lateral Hypothalamus-Caudal | Hypothalamus | 0.2708365 / 0.2610529 (973) | 0.2928438 / 0.2847001 (374) | p = 0.0001388 | d = 0.8520057 |
| Lateral Hypothalamus-Rostral | Hypothalamus | 0.2720674 / 0.2737441 (996) | 0.2799556 / 0.2611548 (323) | p = 0.05259 | d = 0.4334169 |
| Ventromedial Hypothalamus | Hypothalamus | 0.2732628 / 0.2648788 (980) | 0.2863299 / 0.280433 (324) | p = 0.01177 | d = 0.5632742 |
| Periventricular Zone | Hypothalamus | 0.269911 / 0.2585305 (981) | 0.2908163 / 0.2858588 (381) | p = $9.915e^{-05}$ | d = 0.8704249 |
| **Zhang et al., 2014** | | | | | |
| Brainstem | Brainstem | 0.267867 / 0.2592915 (998) | 0.3005153 / 0.2952658 (648) | p = $4.58e^{-12}$ | d = 1.60655 |
| Cerebellum | Cerebellum | 0.270388 / 0.2624734 (912) | 0.3070353 / 0.3015766 (730) | p < $2.2e^{-16}$ | d = 1.879093 |
| Hypothalamus | Hypothalamus | 0.2742063 / 0.2621772 (887) | 0.3043939 / 0.3038627 (649) | p = $3.239e^{-11}$ | d= 1.483688 |

**Supplementary Table 3.** Fisher's exact test was used to compare clock gene ratios in the 90th percentile of positive correlations (FDR < 0.05) from the CCorGsDB by tissue in the mouse and human CNS with the frequency of clock genes found in the mouse and human transcriptomes used as input for WGCNA.

| Transcriptome | # Input genes | # Clock genes | % Clock genes | # CCorGs (90th percentile) | # Recovered Clock genes | % Recovered clock genes | |
|---|---|---|---|---|---|---|---|
| **Mouse** | | | | | | | |
| Isocortex | 19933 | 22 | 0.11 | 1124 | 6 | 0.53 | p = 0.003185* |
| Olfactory Areas | 19933 | 22 | 0.11 | 5162 | 15 | 0.29 | p = 0.006571* |
| Central Nervous System | 19933 | 22 | 0.11 | 1588 | 6 | 0.37 | p = 0.0149* |
| Brain Stem | 19933 | 22 | 0.11 | 2058 | 7 | 0.34 | p = 0.01557* |
| Striatum | 19933 | 22 | 0.11 | 4006 | 11 | 0.27 | p = 0.01765* |
| Thalamus | 19924 | 22 | 0.11 | 2054 | 8 | 0.38 | p =0.04183* |
| Midbrain | 19933 | 22 | 0.11 | 5452 | 9 | 0.16 | p = 0.2816 |
| Pallidum | 19923 | 22 | 0.11 | 3047 | 5 | 0.16 | p = 0.3937 |
| Pons | 19932 | 22 | 0.11 | 1219 | 0 | 0 | p = 0.6352 |
| Hippocampus | 19932 | 22 | 0.11 | 1609 | 2 | 0.12 | p = 0.6994 |
| Hypothalamus | 19885 | 22 | 0.11 | 980 | 1 | 0.10 | p = 1 |
| Cerebellum | 19924 | 22 | 0.11 | 4217 | 4 | 0.11 | p = 1 |
| Medulla | 19932 | 22 | 0.11 | 1662 | 1 | 0.06 | p = 1 |
| **Human** | | | | | | | |
| Central Nervous System | 53921 | 22 | 0.04 | 5015 | 15 | 0.29 | p = 1.367e-07* |
| Cerebellar Hemisphere | 50943 | 22 | 0.04 | 3084 | 10 | 0.32 | p = 7.479e-06* |
| Nucleus Accumbens | 51506 | 22 | 0.04 | 4944 | 12 | 0.24 | p = 1.786e-05* |
| Frontal Cortex | 51004 | 22 | 0.04 | 3038 | 9 | 0.29 | p = 3.671e-05* |
| Caudate | 51311 | 22 | 0.04 | 5415 | 12 | 0.22 | p = 4.221e-05* |
| Spinal Cord | 50064 | 22 | 0.04 | 3822 | 8 | 0.20 | p = 0.0009127* |
| Substantia Nigra | 49596 | 22 | 0.04 | 1858 | 4 | 0.21 | p = 0.01351* |
| Cerebellum | 17526 | 22 | 0.12 | 1015 | 4 | 0.39 | p = 0.05149 |
| Hypothalamus | 51222 | 22 | 0.04 | 1882 | 3 | 0.16 | p = 0.05739 |
| Hippocampus | 17549 | 22 | 0.12 | 1993 | 6 | 0.30 | p = 0.05978 |
| Putamen | 17521 | 22 | 0.12 | 3702 | 9 | 0.24 | p = 0.09748 |
| Amygdala | 17528 | 22 | 0.12 | 2478 | 5 | 0.20 | p = 0.3725 |
| Anterior Cingulate Cortex | 17958 | 22 | 0.12 | 1933 | 3 | 0.15 | p = 0.7301 |

**Supplementary Figure 1.** Top Clock Correlated Genes (CCorGs) by each tissue and the top 3 CCorGs in the integrated Central Nervous System networks, with their respective correlated clock genes in the human data. r = Pearson correlation value.

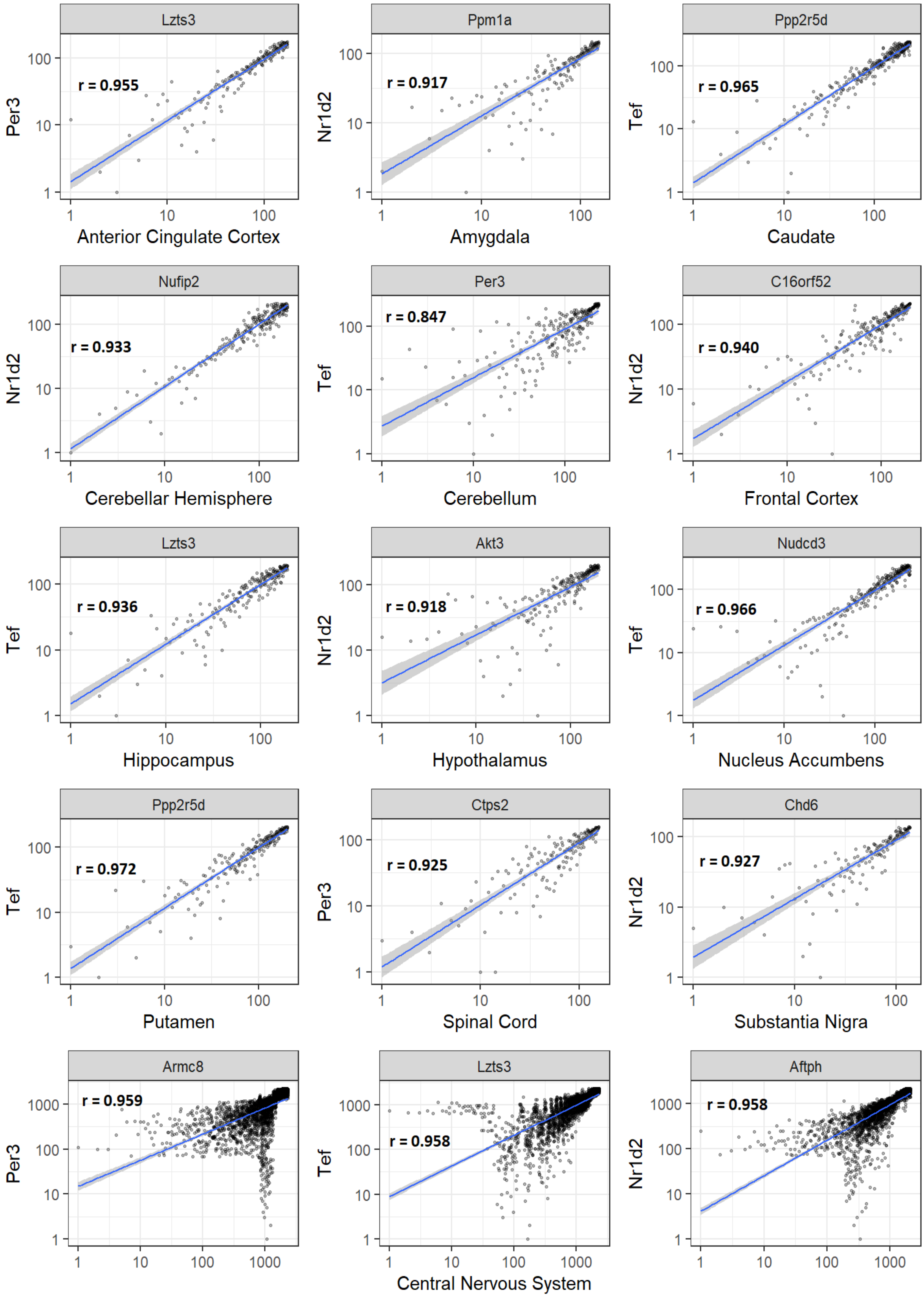

**Supplementary Figure 2.** Top Clock Correlated Genes (CCorGs) by each tissue and the top 3 CCorGs in the integrated Central Nervous System networks, with their respective correlated clock genes in the mouse data. r = Pearson correlation value.

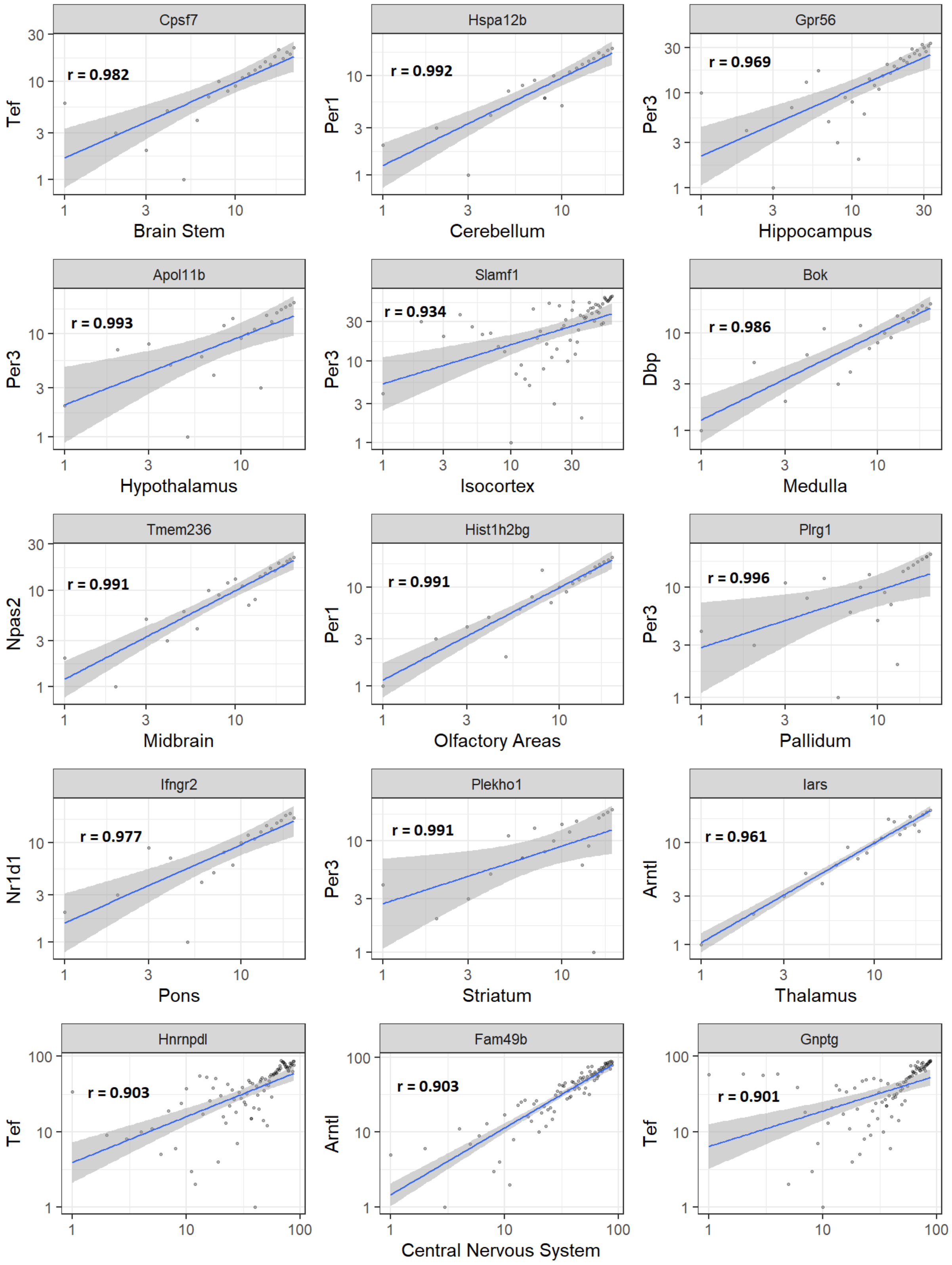

5. LU, Qingqing; KIM, Jin Young. Mammalian circadian networks mediated by the suprachiasmatic nucleus. **The FEBS Journal**, 2021.

6. BEGEMANN, Kimberly; NEUMANN, Anne‐Marie; OSTER, Henrik. Regulation and function of extra‐SCN circadian oscillators in the brain. Acta physiologica, v. 229, n. 1, p. e13446, 2020.

26. HUGHEY, Jacob J.; HASTIE, Trevor; BUTTE, Atul J. ZeitZeiger: supervised learning for high-dimensional data from an oscillatory system. Nucleic acids research, v. 44, n. 8, p. e80-e80, 2016.

34. Allen Institute for Brain Science (2004). Allen Mouse Brain Atlas [dataset]. Available from mouse.brain-map.org.Allen Institute for Brain Science (2011).

40. ZHANG, Bin; HORVATH, Steve. A general framework for weighted gene co-expression network analysis. **Statistical applications in genetics and molecular biology**, v. 4, n. 1, 2005.

46. Piñero, J., Ramírez-Anguita, J. M., Saüch-Pitarch, J., Ronzano, F., Centeno, E., Sanz, F., & Furlong, L. I. (2020). The DisGeNET knowledge platform for disease genomics: 2019 update. Nucleic Acids Research, 48(D1), D845-D855.

47. WISHART, David S. et al. DrugBank: a comprehensive resource for in silico drug discovery and exploration. **Nucleic acids research**, v. 34, n. suppl_1, p. D668-D672, 2006.
